# Utilization of a Human Liver Tissue Chip for Drug-Metabolizing Enzyme Induction Studies of Perpetrator and Victim Drugs

**DOI:** 10.1101/2024.07.17.603946

**Authors:** Shivam Ohri, Paarth Parekh, Lauren Nichols, Shiny Amala Priya Rajan, Murat Cirit

## Abstract

Polypharmacy-related drug-drug interactions (DDIs) are a significant and growing healthcare concern. Increasing number of therapeutic drugs on the market underscores the necessity to accurately assess the new drug combinations during pre-clinical evaluation for DDIs. *In vitro* primary human hepatocyte (PHH)-only models are commonly used only for perpetrator DDI studies due to their rapid loss of metabolic function. But co-culturing non-human cells with human PHHs can stabilize metabolic activity and be utilized for both perpetrator and victim studies, though this raises concerns about human specificity for accurate clinical assessment. In this study, we evaluated Liver Tissue Chip (LTC) with PHH-only liver microphysiological system (MPS) for DDI induction studies. Liver MPS from three individual donors maintained their functional and metabolic activity for up to 4 weeks demonstrating suitability for long-term pharmacokinetics (PK) studies. The responses to rifampicin induction of three PHH donors were assessed using CYP activity and mRNA changes. Additionally, victim PK studies were conducted with midazolam (high clearance) and alprazolam (low clearance) following rifampicin-mediated induction which resulted in a 2-fold and a 2.6-fold increase in midazolam and alprazolam intrinsic clearance values respectively compared to the untreated liver MPS. We also investigated the induction effects of different dosing regimens of the perpetrator drug (rifampicin) on CYP activity levels, showing minimal variation in the intrinsic clearance of the victim drug (midazolam). This study demonstrates the utility of the LTC for *in vitro* liver-specific DDI induction studies, providing a translational experimental system to predict clinical clearance values of both perpetrator and victim drugs.

**SIGNIFICANCE STATEMENT:** This study demonstrates the utility of the Liver Tissue Chip (LTC) with primary human hepatocyte (PHH)-only liver microphysiological system (MPS) for drug-drug interaction (DDI) induction studies. This unique *in vitro* system with continuous recirculation maintains long-term functionality and metabolic activity for up to 4 weeks, enabling the study of perpetrator and victim drug pharmacokinetics, quantification of drug-induced CYP mRNA and activity levels, investigation of patient variability, and ultimately clinical predictions.

## INTRODUCTION

Drug-drug interactions (DDI) can occur when multiple drugs are administered simultaneously, referred to as polypharmacy, leading to adverse drug reactions (ADRs) (Pirmohamed et al. 2004) with profound clinical effects that can reduce therapeutic efficacy or increase drug toxicity. This is a substantial healthcare burden as 37% of older adults use five or more drugs (Gu, Dillon, and Burt 2010) that could potentially lead to DDI-related hospital admissions (Dechanont et al. 2014).

A therapeutic drug may lead to pharmacokinetic (PK) DDI either as a victim, where other drugs affect the investigational drug, or as a perpetrator, where the investigational drug affects concomitant drugs. As most small molecule drugs undergo biotransformation by cytochrome P450 enzymes (CYP) in the liver, CYP-mediated DDIs pose significant risk (Bohnert et al. 2016) and can result in several hundred-fold variations in victim drug exposure in humans (Backman et al. 1998). Thus, the evaluation of DDI risk of new investigational drugs is an essential step prior to clinical first-in-human studies.

Primary human hepatocytes (PHH) are considered the gold-standard for metabolism-based *in vitro* studies; however, a major limitation is the loss of metabolic function over time (Fowler et al. 2020). This limitation challenges long-term drug incubations for perpetrator treatment as well as PK assessments of victim drugs (Lu and Di 2020). To overcome this limitation, multiple *in vitro* co-culture models were developed that cultured PHH with non-human stromal cells or liver non-parenchymal cell lines to extend the culture life of plated PHH (Chan et al. 2019; Hultman et al. 2016). However, the presence of stomal cell-like fibroblasts complicates clearance predictions and can interact with the drugs in a non-physiological manner (Kratochwil et al. 2017). In addition to *in vitro* models, pre-clinical animal models were considered as a viable option for metabolism-related DDI assessments (Prueksaritanont et al. 2006); however, they are rarely used in preclinical DDI studies as differences in species-specific CYP enzyme levels and tissue distribution make animal models less predictive of human enzyme induction (Lu and Li 2001). Therefore, there remains a clear need for a metabolically active and human-specific *in vitro* system for preclinical DDI studies that enables long-term PK assessment of compounds and offers flexibility for testing various dosing regimens.

Human tissue chips, or micro physiological systems (MPS), combine human cells, biomaterials, and biomimetic cues, such as fluid flow, to create physiologically relevant *in vitro* models with sustained metabolic activity. However, current microfluidic-based tissue chips face several challenges, including limited drug exposure in flow-through formats for studying low-clearance drugs, the use of drug-absorbing chip materials, small medium volumes and tissue sizes for assays, and evaporation during long-term PK studies (Fowler et al. 2020). We have addressed these pitfalls with the development of a novel millifluidic system, Liver Tissue Chip (LTC), as described in our previous publication (Rajan et al. 2023). Here, we evaluated the LTC system for *in vitro* induction studies for multiple PHH donors. The LTC system was further evaluated for CYP-mediated DDI studies by inducing the system with rifampicin, a perpetrator drug, and conducting victim drug PK assessments with the known clinical victim drugs, midazolam and alprazolam.

## METHODS

### Javelin’s LTC System

The LTC system from Javelin Biotech, Inc. used in this study includes gamma-sterilized chips (Fig. 1A) and multiplex controllers (Fig. 1B). The basic working principles were described previously by Rajan et al. 2023. Briefly, the thermoplastic-based LTC houses a 1-cm^2^ cell culture chamber with a removable sealing lid that allows for direct seeding into the chamber and access to the culture throughout the study. The on-board pump, controlled by the multi-plex controller, allows for medium recirculation during the study at a set flow rate of 2 mL/h. The oxygenation chamber allows for continuous oxygen diffusion into the medium through a gas-permeable membrane, providing a constant oxygen supply to the cells. This chamber also allows for aseptic medium sampling for multiple kinetic timepoints during long-term PK studies.

**Figure 1:**
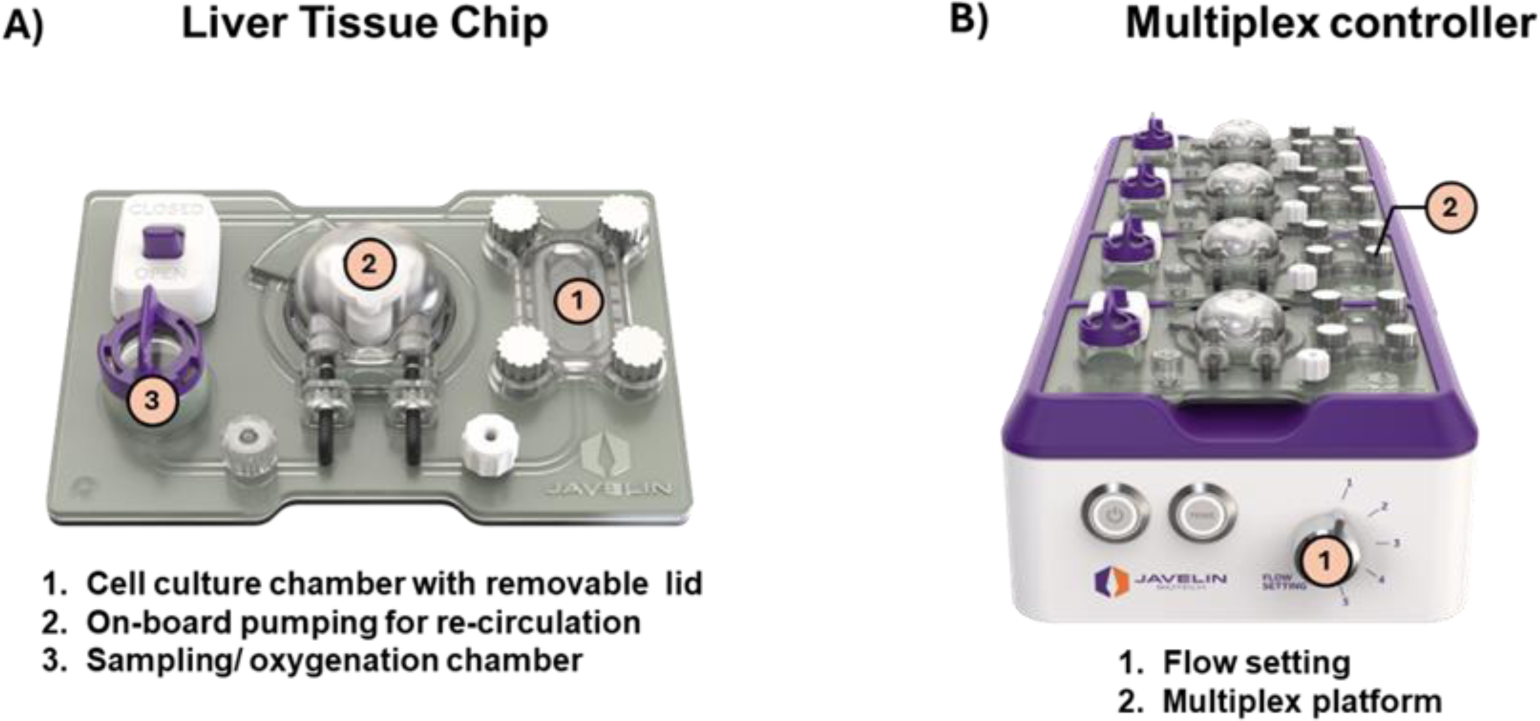
Human liver tissue chip (LTC) system. The LTC system includes (A) LTC with key design features that enables long term hepatic culture and (B) Multiplex controller that controls the medium recirculation of up to 4 connected LTCs at predetermined flow rate.

### Liver MPS Culture and Maintenance

#### PHH Culture

The liver MPS was comprised of PHH in sandwich culture format within the LTC culture chamber. Before cell seeding, the culture chamber was coated with 100 µg/mL collagen (Corning, Corning NY) and 25 µg/mL fibronectin (Sigma Aldrich, St Louis MO) prepared in phosphate-buffered saline followed by overnight incubation at 4⁰C and 2-h incubation at 37°C.

Cryopreserved PHHs were purchased from LifeNet Health, Virginia Beach VA (Donor 2214423, Female), AnaBios Corporation, San Diego CA (Donor 1045, Female), and ThermoFisher Scientific, Waltham MA (Donor Hu8339, Female). Donor information including age, ethnicity, and cause of death is provided in Table S6. Cryopreserved PHH vials were thawed at 37⁰C for 2 min and the cell suspension was transferred to pre-warmed hepatocyte thawing medium (Gibco, Waltham MA). This suspension was centrifuged at 100 × *g* for 8 min and then re-suspended in hepatocyte plating medium (Gibco, Waltham MA) at a density of 0.86 million cells/mL. Each cell chamber was seeded with 215,000 cells and incubated overnight. The attached cells were overlayed with 0.35 mg/mL Matrigel (Corning, Corning NY) prepared in ice-cold hepatocyte maintenance medium (Gibco, Waltham MA). Following overnight incubation, the cell chamber was sealed with the lid, and the chip was filled with 1.8 mL of maintenance medium and connected to flow for culture maintenance.

#### Liver MPS Maintenance

The liver MPS was maintained by replacing the spent medium every 4–7 days. The collected spent medium was used to run multiple biochemical assays quantifying albumin and urea production. The hepatocyte morphology was monitored every two days using bright-field microscopy (Zeiss).

### Functional and Metabolic Activity Assessment

Albumin production was quantified by R-plex human albumin assay (Meso Scale Discovery, Rockville MD) and urea synthesis was measured by colorimetric assay (BioAssay Systems, Hayward CA). Albumin and urea values were adjusted for total media volume and number of days elapsed between media changes and reported as µg/day/million hepatocytes.

The liver MPS was incubated with a probe substrates cocktail for five major CYP isoforms (3 µM midazolam, CYP3A4; 90 µM diclofenac, CYP2C9; 3 µM R-omeprazole, CYP2C19; 3 µM tacrine, CYP1A2; 20 µM dextromethorphan, CYP2D6) for 3 h under recirculation at 37°C and metabolism was assessed by metabolite quantification of 1-OH midazolam, 4-OH diclofenac, 5-OH omeprazole, OH-tacrine, and dextrorphan, respectively using LC-MS/MS (see Supplementary Information).

### Quantitative Real-Time PCR (qPCR)

For qPCR analysis, RNA was extracted from the liver MPS using a guanidine thiocyanate-containing lysis buffer (Invitrogen, Waltham MA) and further purified per the vendor protocol (Invitrogen, Waltham MA). The total extracted RNA was converted to cDNA (Applied Biosystems, Waltham MA), loaded onto custom Taqman array plates (ThermoFisher, Waltham MA), and run with the StepOnePlus Real-Time PCR system (ThermoFisher, Waltham MA). The fluorescence emitted during the amplification process was measured at each cycle, and the cycle threshold (Ct) values were determined. Relative gene expression levels were calculated using the 2^(−ΔΔCt) method with normalization to two reference genes, 18S and ACTB. The ΔCt was calculated by subtracting the Ct value from the control samples from the Ct value of the experimental group, and ΔΔCt was obtained by comparing the ΔCt of each target gene to that of the reference genes.

### Liver MPS Induction using Rifampicin

The Liver MPS was dosed with rifampicin for 72 h to investigate the effects of three rifampicin dosing regimens: i) daily spiking, ii) daily fresh dose, and iii) single dose. On day 3, using the complete medium change protocol, 1.8 mL of fresh medium containing 10 µM rifampicin was added for all three-dosing regimens. For the daily spiking regimen, the medium in the sampling chamber was spiked with 1.8 µL of 10 mM rifampicin stock every 24 h after the initial dose for an effective concentration of at least 10 µM. In the daily fresh dose condition, spent rifampicin-dosed medium was replaced every 24 h with fresh medium containing 10 µM rifampicin until 72 h. For the single dose regimen, no drug spiking or medium replacement was conducted throughout the 72 h of induction pre-treatment. At the end of rifampicin incubation, metabolic activity and/or mRNA changes in the liver MPS were evaluated using probe substrate cocktail assay and qPCR analysis, respectively.

### Long-term PK Assessment with Victim Drugs

Long-term PK of the victim drugs, midazolam and alprazolam, was assessed after treatment with the perpetrator drug, rifampicin. The substrates midazolam (1 µM) or alprazolam (1 µM) were added to the liver MPS as a bolus dose on day 6 in the presence and absence of 10 µM rifampicin. Medium samples of 50 µL were collected from the sampling chamber every 24 h for 72 and 120 h for midazolam and alprazolam, respectively. Samples were analyzed for parent drug depletion and metabolite formation using LC-MS/MS (see Supplementary Information). Area-under-the curve (AUC) of the substrate (victim drug) depletion curve was calculated using GraphPad Prism and was represented as AUC _0-t_ where ‘t’ represents the endpoint of victim drug study. *In vitro* intrinsic clearance (CL_int_ [µL/min/10^6^ hepatocytes]) was calculated using the slope of substrate depletion data and predicted in vivo intrinsic clearance (CL_h_ [mL/min/kg]) was estimated after physiological scaling of human factors. Equations and experimental values used for scaling *in vitro* parameters are previously described in Rajan et al. 2023. Briefly, unbound *in vitro* intrinsic clearance (CL_int(u)_) was calculated by dividing CL_int_ by fraction unbound in the media (fu_media_) which was scaled to human liver equivalent unbound intrinsic clearance (CL_int(u), H_ [mL/min/kg]) taking into account human hepatocellularity and liver weight. Finally, the upscaled human CL_int(u)_ was converted to predicted human hepatic clearance (CL_h_ [mL/min/kg]) using well-stirred (WS) and parallel tube (PT) models.

### Drug Stock Preparations

Drugs were purchased in powdered form from Sigma-Aldrich, St Louis MO (rifampicin, diclofenac, dextromethorphan), Cayman Chemicals, Ann Arbor MI (R-omeprazole), and Tocris, Bristol UK (tacrine). Drugs were dissolved in dimethyl sulfoxide (DMSO) to prepare stock solutions at a concentration at least 1000-times higher than the required final concentration; stock solutions were stored at –80°C until use. Midazolam and alprazolam (Sigma-Aldrich, St Louis MO) were purchased pre-solubilized in methanol; 1 mM stock solutions were prepared in DMSO and diluted to final concentration of 1 µM in medium. All drug incubations in the LTC were conducted with a DMSO vol/vol percentage of <0.5.

### Statistical Analysis

To determine statistical significance, unequal variance t-tests were performed between two conditions and the level of significance was determined using p-value ranges, with p<0.05 deemed significant. A minimum of three independent biological replicates were performed for each experimental group. Microsoft Excel and GraphPad Prism were used for data analysis and GraphPad Prism was used to plot the data.

## RESULTS

### Long-term Maintenance of Liver MPS from Multiple PHH Donors

In the LTC system, PHH from three different donors (2214423, 1045, and Hu8339) were maintained in sandwich culture format for over 4 weeks using the optimized protocol (Fig. 2A). The functionality of the liver MPS was evaluated by measuring the production of albumin and urea, while monitoring PHH morphology using bright field imaging. PHH from all three donors maintained their cobblestone morphology (Fig. 2B and Fig S1) and prolonged production of albumin and urea throughout the culture period of 4-weeks (Fig. 2C). Healthy liver MPS produced an average albumin level per day ranging from ∼20–50 µg/day/10^6^ hepatocytes during the culture duration, which is within physiological levels (Baudy et al. 2020). The PHH culture from Hu8339 showed donor variability toward the end of the culture period, where albumin levels were below 20 µg/day/10^6^ hepatocytes after day 15. All donors displayed higher levels of urea production on day 3 of culture followed by a sustained production with average levels ranging from ∼100–300 µg/day/10^6^ hepatocytes. The stable production of albumin, a protein synthesis marker, and urea, a metabolic function marker, demonstrates the maintenance of long-term healthy liver MPS with sustained hepatocyte phenotype and viability.

**Figure 2:**
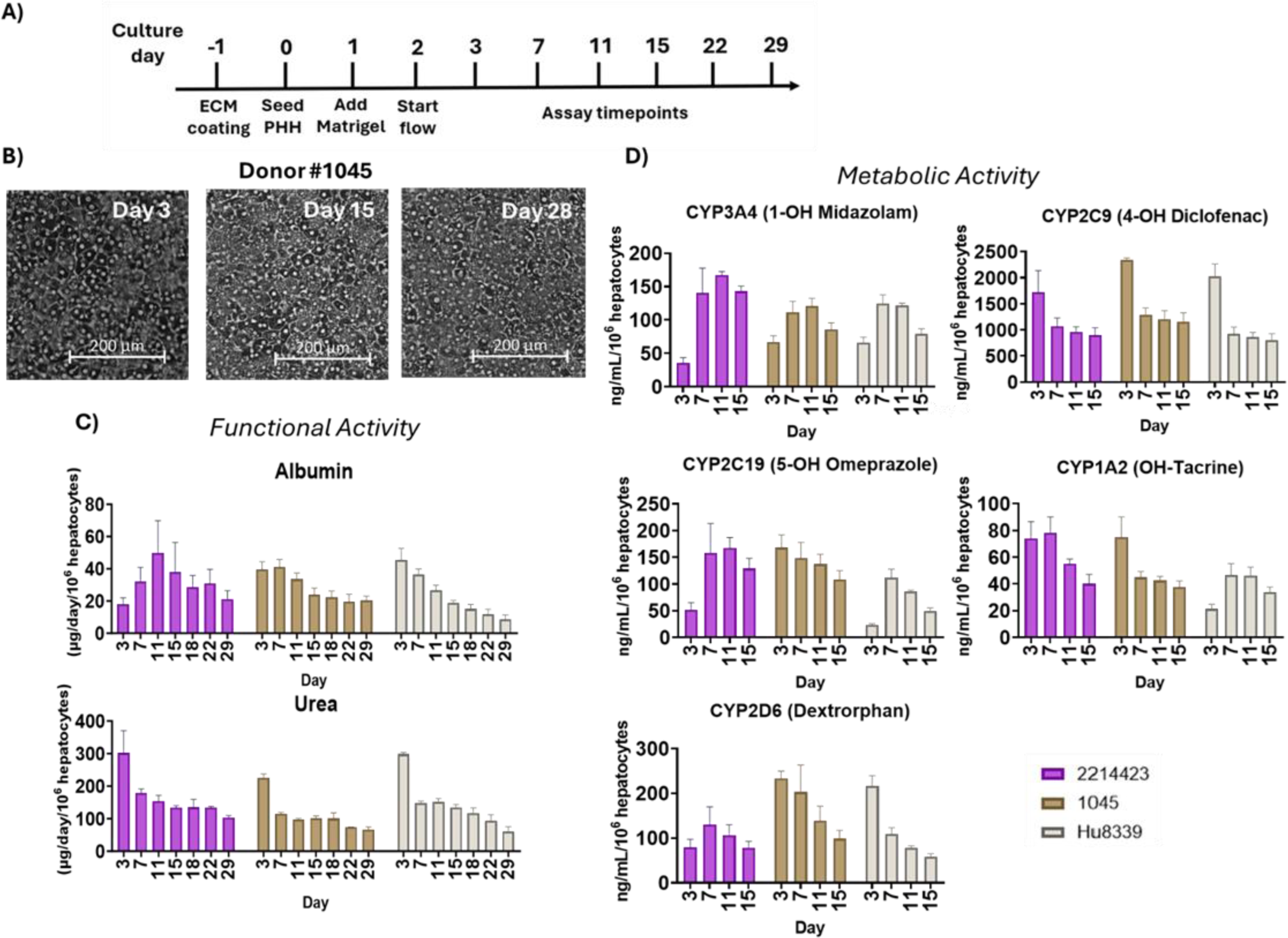
Biological characterization of PHHs from three donors (2214423, 1045, Hu8339) maintained in the LTC. A) Optimized experimental workflow for long-term liver MPS maintenance in LTC. B) Brightfield images of PHH (Donor 1045) showing tissue morphology that was maintained for at least 28 days. Liver MPS from all three donors sustained long-term (C) functional activity quantified by measuring albumin and urea production until day 29 and (D) metabolic activity measured by incubating the liver MPS with 5-in-1 probe substrates cocktail for CYP3A4, CYP2C9, CYP2C19, CYP1A2, and CYP2D6 and quantifying associated metabolites until at least day 15. Plotted data represents mean + SD of 3-4 biological replicates.

PHHs from all the donors sustained metabolic potential for at least 15 days, confirming metabolically active and stable liver MPS for DDI studies (Fig 2D). CYP3A4 and CYP2C19 activity levels (measured by quantifying 1-OH midazolam and 5-OH omeprazole, respectively) showed an increase for all three donors from day 3 of the culture period, with sustained activity from day 7 except for CYP2C19 activity of 1045, which showed sustained activity levels from day 3 onward. In contrast, activity levels of CYP1A2, CYP2C9, CYP2D6 (quantified by OH-tacrine, 4-OH diclofenac, and dextrorphan, respectively) showed higher activity on day 3, which is a day after flow, followed by sustained activity during the culture period. Donor variability was observed for CYP1A2 activity levels in Hu8339 and CYP2D6 activity levels in 2214423, where an increase in enzyme activity levels was observed after day 3 for both the donors, with activity levels stable in later days of culture. This demonstrates the ability of LTC to preserve the long-term CYP activity of PHH from multiple donors while capturing their inter-donor differences, making it a well-suited *in vitro* platform for conducting metabolic studies and achieving more accurate predictions.

### Effects of Different Rifampicin Dosing Regimens on CYP Activity Levels

The liver MPS (donor 1045) was induced with rifampicin using three dosing regimens over 72 h: (i) daily spiking, (ii) daily fresh dose, and (iii) single dose (Fig. 3A and Fig. S2A). All three rifampicin dosing regimens showed similar levels of induced CYP3A4, CYP2C19, and CYP1A2 activities in LTC. Interestingly, the daily fresh regimen resulted in significantly higher CYP2C9 activity than the other two regimens (Fig. 3B). For each dosing regimen, the rifampicin concentration in LTC was quantified daily during the induction period between days 4 and 6 (Fig. S2B–D). Rifampicin exposure was calculated by quantifying the AUC of the concentration over 72 h of induction pre-treatment. The AUC results revealed the highest rifampicin exposure of 22.4 µM*day for the daily spiking regimen, 15.9 µM*day with the fresh daily dose, and the lowest exposure of 6.6 µM*day for the single rifampicin dose (Fig. 3C). Based on the highest rifampicin exposure, the daily spiking regimen was used for subsequent studies.

**Figure 3:**
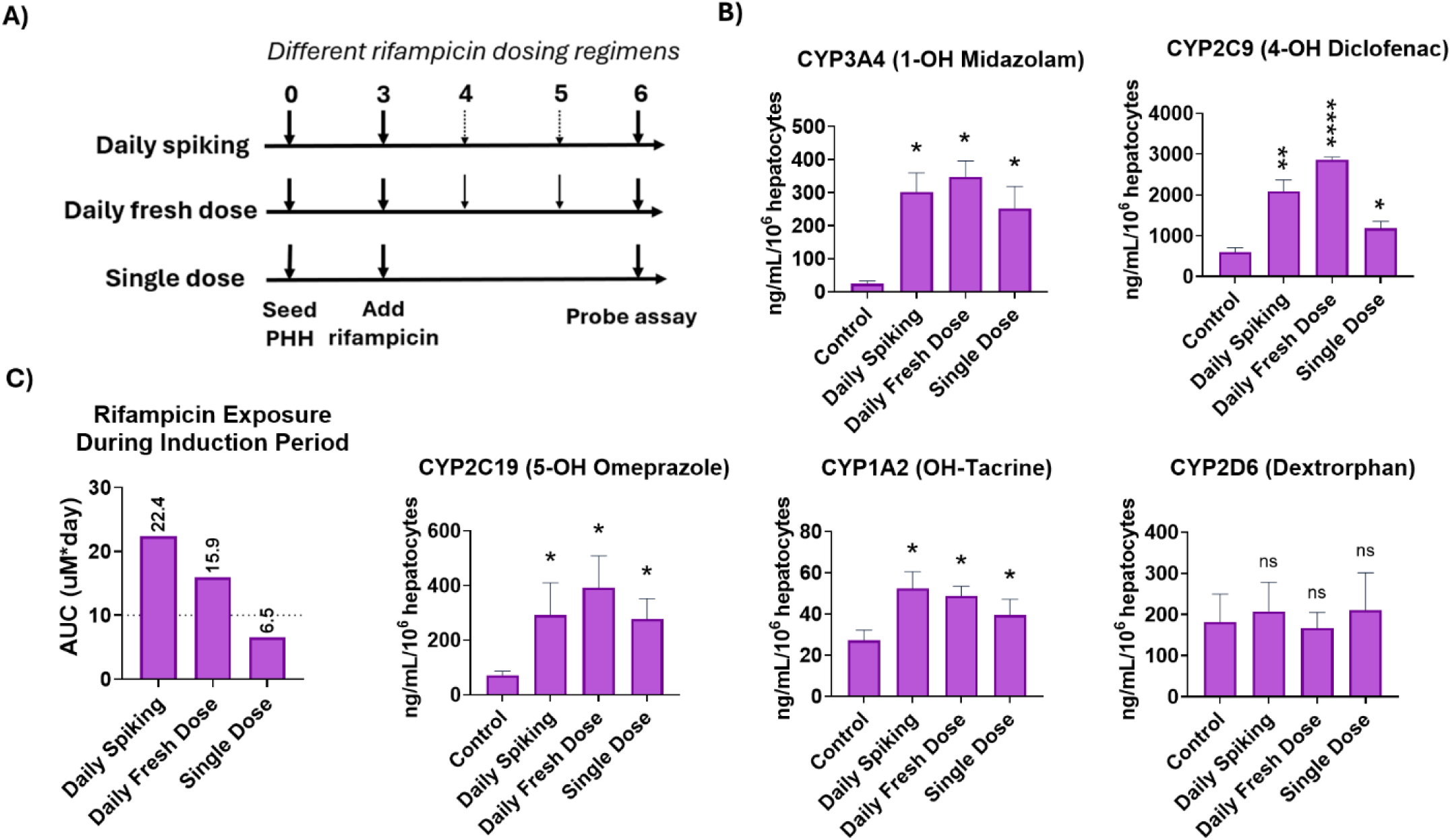
Investigating induction effects of rifampicin on LTC across different rifampicin dosing regimens (PHH donor: 1045) (A) Experimental workflow of dosing regimens: daily spiking, daily fresh dose, and single dose. (B) Activity levels of CYP3A4, CYP2C9, CYP2C19, CYP1A2, and CYP2D6 enzymes were quantified using 5-in-1 probe substrate cocktail assay after rifampicin treatment across different dosing regimens. Compared to the vehicle control, the liver MPS showed dose regimen-dependent variation only for CYP2C9 levels and no significant differences among regimens for other measured CYP enzymes (significance level not shown). (C) shows the rifampicin exposure to liver MPS for different dosing regimens represented as AUC of rifampicin concentration-time curve obtained during three days of induction pre-treatment period. Statistical significance is displayed relative to control determined using un-equal variance t-test (* .001 < p < 0.05; ** .0001 < p < .001; *** .00001 < p < .0001; **** p < .00001; no significance (ns) p > 0.05). Plotted data represents mean + SD of 3-4 biological replicates.

### Evaluation of Donor Variability in Rifampicin-induced CYP mRNA Expression and Activity Levels

Liver MPS from three donors were induced with rifampicin on day 3 for 72 h using the daily spiking regimen (Fig. 4A). At the end of the induction period, CYP enzymatic activity was assessed using a 5-plex CYP probe cocktail and CYP mRNA levels were assessed by qPCR. Rifampicin treatment induced CYP3A4, CYP2C9, and CYP2C19 activities for all donors compared to the vehicle control (Fig. 4B & C). While CYP2D6 activity was not affected by rifampicin treatment with any of the donors, CYP1A2 induction was observed with one donor, 1045. CYP3A4 activity was induced 1.3-, 1.8-, and 2.7-fold for 2214423, 1045, and Hu8339, respectively, while fold-increase of mRNA expression levels (from 2.3 to 7.6-fold increase) was higher than enzymatic activity levels from each donor (Fig. 4C). Interestingly, CYP2C9 mRNA expression after induction was <2-fold for each donor; however, the range of increase in the enzymatic activity was higher (1.8-to 5.8-fold) than the mRNA fold increase range (from 0.63 to 1.6-fold).

**Figure 4:**
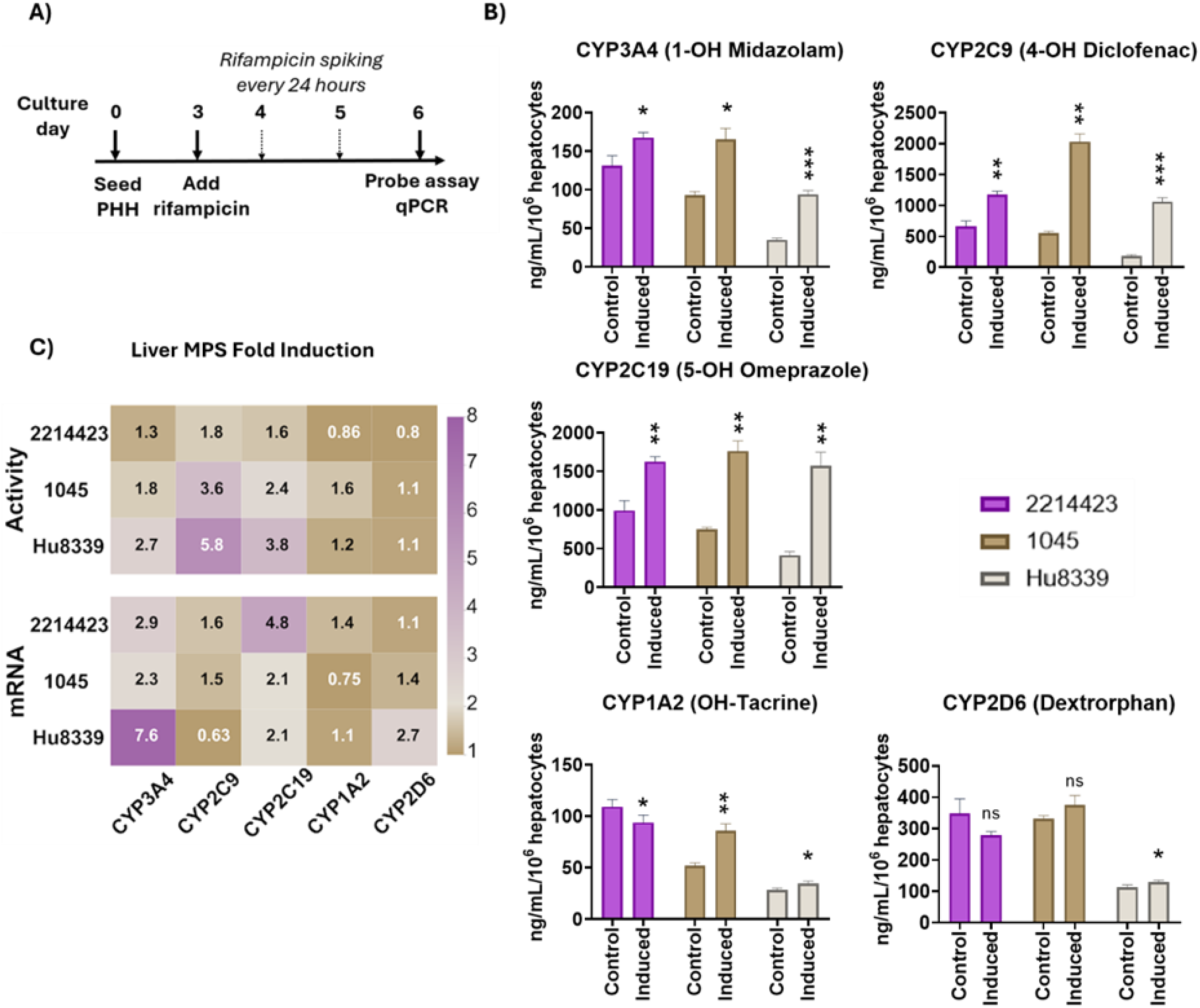
Comparison of rifampicin-mediated induction effects on Liver MPS across different donors (2214423, 1045, Hu8339) (A) Experimental workflow with optimized daily spiking dosing regimen. (B) Enzyme activity levels of CYP3A4, CYP2C9, CYP2C19, CYP1A2, and CYP2D6 quantified using 5-in-1 probe substrate cocktail assay shows rifampicin-mediated induction compared to the vehicle control for all three donors (C) Fold induction changes for mRNA and enzyme activity levels across different donors demonstrate donor variability observed in induced levels of enzyme activity from control. Statistical significance is displayed relative to control determined using an un-equal variance t test (* .001 < p < 0.05; ** .0001 < p < .001; *** .00001 < p < .0001; **** p < .00001; no significance (ns) p > 0.05). Plotted data represents mean + SD of 3-4 biological replicates.

### Baseline Recovery of CYP Activity Levels After Rifampicin Withdrawal

The liver MPS (donor 1045) was induced with rifampicin starting at day 3 for 72 h using the daily spiking regimen (induction pre-treatment period); at day 6, a single dose of 10 µM rifampicin was added for another 72 h (victim PK study period). On day 9, rifampicin-free hepatocyte maintenance medium was added for an additional 6 days to observe the baseline recovery of CYP activity levels (Fig. 5A and Fig. S3A). CYP activity levels were assessed on days 6, 9, 12, and 15. After the induction pre-treatment period (day 6) and at the end of victim PK study period (day 9), CYP3A4, CYP2C9, and CYP2C19 activities were maintained at the high levels, compared to the vehicle control, expected of an induced condition. After removing rifampicin on day 9, CYP3A4- and CYP2C19-induced activity returned to baseline levels within 3 days and CYP2C9 reduced within 6 days (Fig. 5B). Additionally, we characterized the effects of the daily fresh dose and single dose regimens of rifampicin on CYP activity recovery. The single dose regimen of rifampicin resulted in similar reduction kinetics of CYP3A4 and CYP2C19 activity as daily spiking. However, CYP2C9 activity returned to baseline levels within 3 days rather than the 6 days observed after daily spiking of rifampicin. The daily fresh dose regimen also resulted in similar CYP3A4 activity recovery as daily spiking; however, CYP2C9 and CYP2C19 activity did not completely return to baseline levels after induction pre-treatment (Fig. S3B).

**Figure 5:**
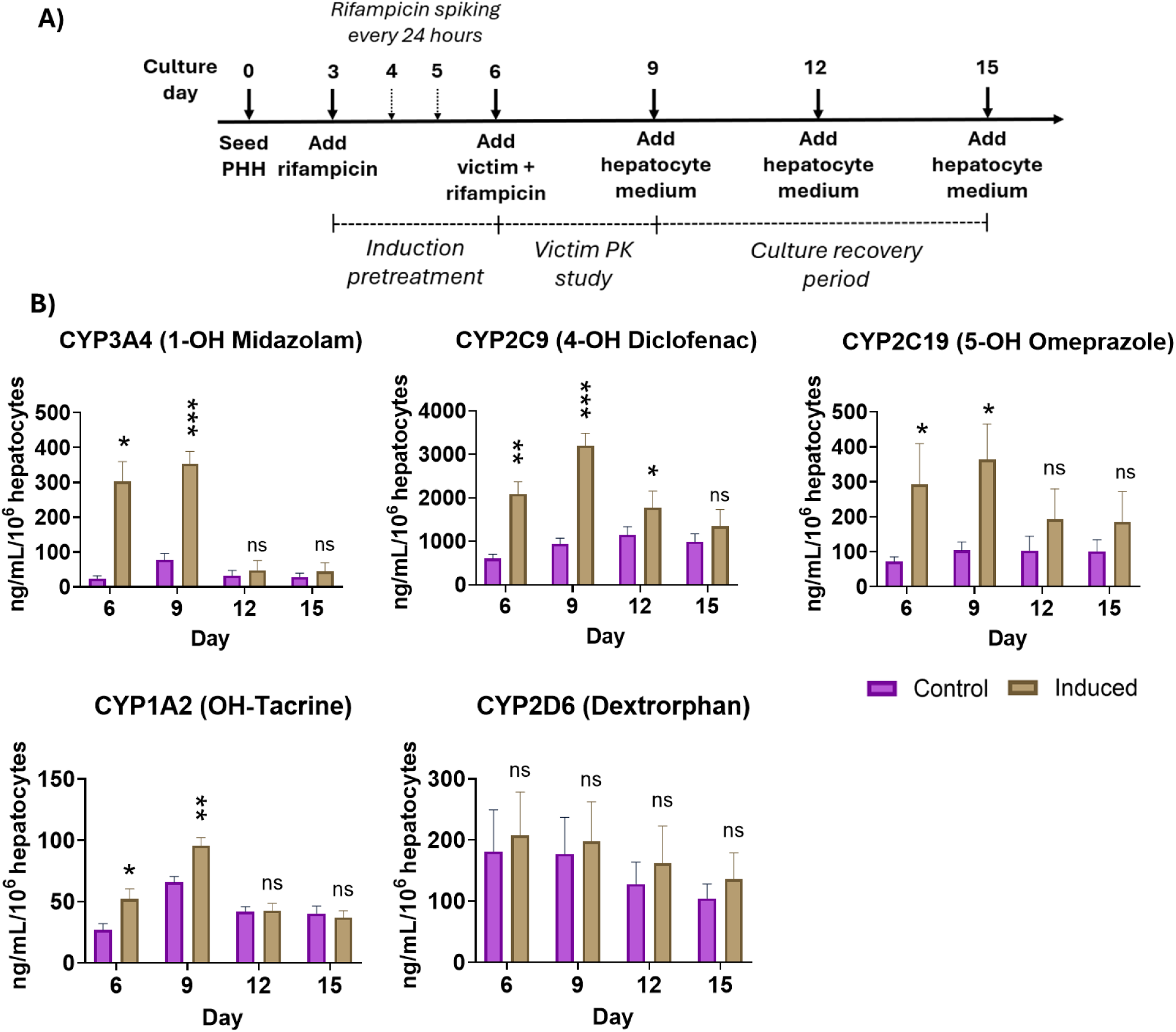
Time course and enzyme activity levels of liver MPS during long-term DDI study involving rifampicin mediated induction, victim PK study, and de-induction after cessation of rifampicin (PHH donor: 1045). (A) Experimental workflow of DDI victim-perpetrator study with recovery time course. (B) Time course plots of CYP activity levels (CP3A4, CYP2C9, CYP2C19, CYP1A2, and CYP2D6) shows increase in activity during rifampicin treatment (day 3 – day 9) compared to the vehicle control followed by de-induction that is observed by activity levels returning to baseline levels after cessation of rifampicin treatment on day 9. Statistical significance is displayed relative to control determined using un-equal variance t-test (* .001 < p < 0.05; ** .0001 < p < .001; *** .00001 < p < .0001; **** p < .00001; no significance (ns) p > 0.05). Plotted data represents mean + SD of 3-4 biological replicates.

### Victim Drug Intrinsic Clearance Changes in Response to a Perpetrator

The liver MPS (donor 1045) was induced with rifampicin starting at day 3 for 72 h; then, at day 9, 1 µM victim drugs (midazolam or alprazolam) were co-dosed with 10 µM rifampicin, the perpetrator drug, for victim PK studies (Fig. 6A and Fig. S4A). Midazolam, a high clearance compound, and alprazolam, a low clearance compound, are both mostly metabolized by CYP3A4 enzyme. We observed higher intrinsic clearance values for each drug in the rifampicin induction group compared to the untreated liver MPS. The midazolam victim study showed a reduction of 43% in the AUC_0 - 72hr_ of parent drug concentration compared to the control group (Fig. 6B). Unbound *in vitro* intrinsic clearance for the rifampicin-induced liver MPS (19.14 µL/min/million cells) and vehicle control (9.51 µL/min/million cells) showed a 2-fold increase in midazolam clearance (Fig. 6E). The predicted human hepatic clearance values of midazolam using the well-stirred model and the parallel tube model were 4.23 mL/min/kg and 4.67 mL/min/kg for the induced liver MPS and 2.34 mL/min/kg and 2.47 mL/min/kg for the control, respectively (Table 1). Daily fresh and single dose regimens of rifampicin induction resulted in 54% and 42% reduction in the AUC_0 – 72hr_ of the midazolam depletion curve, with 2.2- and 1.98-fold increase in intrinsic clearance values, respectively (Fig. S4B & C), resulting in no significant differences between any of the three dosing regimens. The primary metabolite of midazolam (1-OH Midazolam) was quantified to demonstrate inducibility and increased metabolite production in the presence of rifampicin. There was a 2.1-fold increase in the AUC of 1-OH midazolam formation rate from rifampicin-induced liver MPS compared to the vehicle control (Fig 6D). For alprazolam, the AUC_0 – 120hr_ of the parent drug was reduced by 2ic 2% for the induced liver MPS (Fig. 6C), while the unbound intrinsic clearance values were increased from 0.82 µL/min/million cells for vehicle control to 2.13 µL/min/million cells for induced liver MPS, resulting in a 2.6-fold increase (Fig 6E). The predicted human hepatic clearance estimates of alprazolam was also calculated using the well-stirred model and the parallel tube model with resulting values of 2.33 mL/min/kg and 2.46 mL/min/kg for the induced liver MPS and 0.96 mL/min/kg and 0.98 mL/min/kg for the vehicle control, respectively (Table 1)

**Figure 6:**
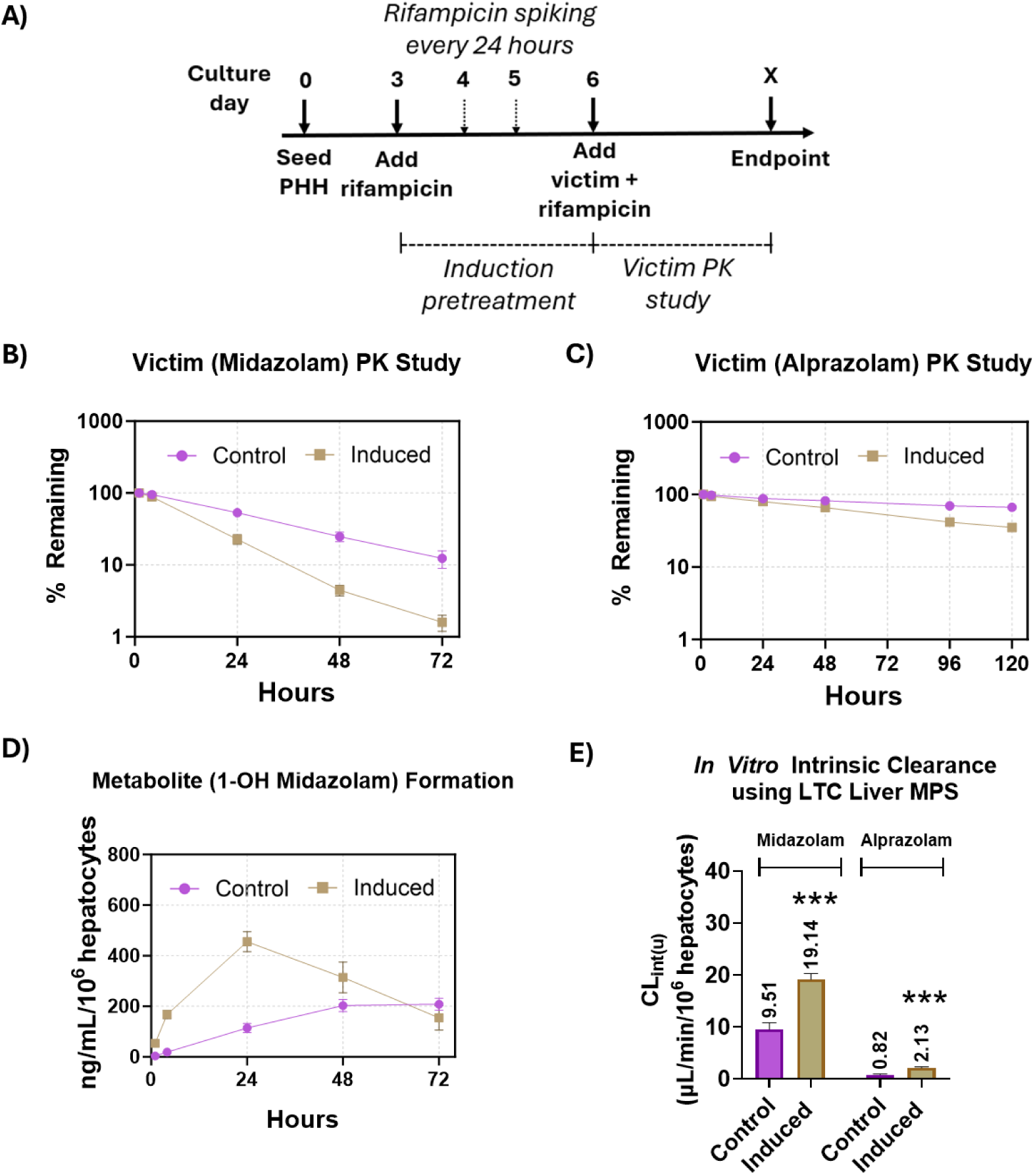
Drug-Drug Interaction (DDI) observed in LTC during long-term victim-perpetrator study (PHH donor: 1045). (A) Experimental workflow of victim-perpetrator PK study with daily spiking of rifampicin regimen during induction pre-treatment. The PK profile of both (B) high-turnover drug midazolam and (C) low-turnover drug alprazolam shows increased depletion rate in rifampicin induced liver MPS compared to the vehicle control. (D) shows increased production of primary metabolite (1-OH midazolam) of midazolam in induced liver MPS compared to the control. (E) Unbound in vitro intrinsic clearance of victim drugs from induced and control liver MPS. Statistical significance is displayed relative to control determined using un-equal variance t-test (* .001 < p < 0.05; ** .0001 < p < .001; *** .00001 < p < .0001; **** p < .00001; no significance (ns) p > 0.05). Plotted data represents mean + SD of 3-4 biological replicates

**Table 1.** Predicted Human Hepatic Clearance (CL_h_) Values Using Well Stirred (WS) and Parallel Tube (PT) Model Unbound *in vitro* intrinsic clearances (CL_int(u)_) of the victim drugs (midazolam and alprazolam) were calculated for both rifampicin induced and vehicle control using the LTC substrate depletion data which was scaled to predicted human hepatic clearance values using human hepatocellularity and WS, PT models.

| Drug | Condition | $CL_{int(u)}$<br>[$\mu\text{L}/\text{min}/10^6$ hepatocytes] | $CL_h$ (WS)<br>[mL/min/kg] | $CL_h$ (PT)<br>[mL/min/kg] |
| --- | --- | --- | --- | --- |
| Midazolam | Control | 9.51 | 2.34 | 2.47 |
|  | Induced | 19.14 | 4.23 | 4.67 |
| Alprazolam | Control | 0.82 | 0.96 | 0.98 |
|  | Induced | 2.13 | 2.33 | 2.46 |

## DISCUSSION

Prediction of CYP-mediated DDI risk of a new investigational drug during the pre-clinical drug development process is an essential step before clinical studies. Such assessment requires an *in vitro* system capable of maintaining long-term hepatic metabolic activity of PHHs to conduct intrinsic clearance estimation and enzyme induction studies (Fowler et al. 2020). To demonstrate the utility of LTC for *in vitro* DDI studies, we evaluated the system for long-term functional and metabolic activities of multiple PHH donors and performed rifampicin induction of multiple CYP isoforms to obtain clinically comparable results for victim drug intrinsic clearance changes in response to a perpetrator drug.

LTC maintained prolonged functional activity of healthy PHHs from three different donors, producing physiologically relevant albumin levels. The observed albumin production range of 20–50 µg/day/10^6^ hepatocytes is comparable to the existing long-term complex micropatterned co-culture model consisting of PHHs and non-human stromal cells (Khetani and Bhatia 2008), showing the novelty of the engineered PHH-only millifluidic *in vitro* system in extending the functionality of plated PHHs long-term. All donors sustained long-term metabolic activity, as demonstrated by multiple CYP isoforms. These results indicated the applicability of LTC system for the long-term PK studies. CYP1A2, CYP2D6, and CYP2C9 activity profiles showed similar trends of higher activity 1-day post introduction of flow-induced shear stress and sustained expression throughout, as opposed to CYP3A4 and CYP2C19 activity levels where an increase in CYP activity was observed as the culture progressed. This could be due to the varying effects of fluidic flow on different CYP isoforms (Du et al. 2017)

Donor variability was observed for multiple CYP isoforms, which displayed different activity levels and profiles. Such variability in metabolic activity is also observed in the clinic and depends on multiple factors including sex, age, and ethnicity, which can all impact clinical outcomes (Zanger and Schwab 2013). Higher clearance of midazolam has been reported in South Asian compared to Caucasian populations (van Dyk et al. 2018), and women exhibited higher clearance of the CYP3A substrate, midazolam (Zhu et al. 2003). Thus, PHH-based liver MPS can be used as an *in vitro* research tool to study individual donors from various backgrounds to predict clinical profiles for special populations.

The inducibility of PHH from multiple donors was assessed by measuring both CYP mRNA and activity levels. We observed more consistent induction of mRNA levels than activity except for with CYP2C9, suggesting that mRNA is a more sensitive induction marker than activity (Fahmi et al. 2010). All three donors were induced >2-fold for CYP3A4 and CYP2C19 mRNA, which is above the threshold of a positive *in vitro* induction signal from treatment with rifampicin (Kenny et al. 2018). CYP3A4 activity fold induction levels did not reach more than 2-fold for two donors, which could be due to high baseline activity levels of the donors indicating a stable and metabolically active healthy liver MPS. Nevertheless, CYP3A4, CYP2C9, and CYP2C19 activity levels of all three hepatocyte donors were increased significantly in response to rifampicin treatment. We also studied the de-induction of CYP activity levels in the liver MPS after rifampicin withdrawal and compared it to the baseline control group. CYP3A4 activity levels returned to baseline within 3 days after being in the presence of rifampicin for 6 days, including induction pre-treatment with multiple dosing regimens and the victim PK study conducted in the presence of rifampicin. Such *in vitro* studies monitoring long term metabolic activity levels could provide insight for clinical CYP activity recovery after being induced, impacting victim drugs and potentially helping improve clinical trial designs (Imai et al. 2008; Reitman et al. 2011)

Different dosing regimens - daily spiking, daily fresh, and single doses - of the perpetrator drug, rifampicin were investigated for their CYP induction potential. Daily spiking resulted in the highest exposure of rifampicin followed by daily fresh dose and then single dose. Despite these different exposures, CYP3A4 activity was induced to the same levels with all dosing regimens, leading to no significant changes in midazolam intrinsic clearance compared to vehicle control. Such studies could provide insight into dosing regimens of perpetrator drugs and associated DDI, potentially incorporating multiple scenarios and enabling the selection of appropriate dosing strategies before clinical studies are conducted (Herman et al. 1983; Xu et al. 2011).

*In vitro* assessment of victim drug intrinsic clearance after induction is a challenge, especially for low-clearance compounds as longer incubation times with metabolically active liver cultures are required (Di and Obach 2015). Here, we demonstrated victim PK studies with both a low hepatic clearance compound, alprazolam, and a high hepatic clearance compound, midazolam. The intrinsic clearance of midazolam in the LTC increased 2-fold after induction and the AUC_0 – 72hr_ of midazolam depletion curve reduced by 43%. This is comparable to observed clinical results, where the AUC_0-∞_ of intravenously dosed midazolam was reduced by 55%, leading to a 2.18-fold increase in systemic clearance after rifampicin administration for 7 days (Gorski 2003). Similar to midazolam, there was a 2.6-fold increase in alprazolam intrinsic clearance in rifampicin-induced LTC, with a reduction of 22% in AUC_0 – 120hr_ of alprazolam. In addition to long-term metabolically stable liver MPS, continuous recirculation in the LTC without any medium change during the victim PK study allowed for the accumulation of primary metabolite, 1-OH midazolam, generated from parent drug, midazolam. Both parent drug and primary metabolite were quantified from the multi-timepoint collected samples for both induced and control LTC, displaying higher metabolite generation after induction. Such a design could also be applied to studies where the effects of generated metabolites need to be evaluated pre-clinically to avoid unexpected interactions in the clinic (Steinbronn et al. 2021; Templeton et al. 2016).

In this study, we focused on induction-based DDIs occurring at the hepatic level using a liver-only MPS system that supports the assessment of intravenously dosed drugs. However, oral administration is the most common drug administration route, and drugs having high intestinal metabolism are more susceptible to inducer-based DDI (Xie, Ding, and Zhang 2016). For example, midazolam, which is mostly metabolized by CYP3A4, is metabolized almost equally in the liver and gut. Clinically, this leads to an AUC change of 98.5% in the plasma concentration-time curve of orally dosed midazolam after rifampicin treatment compared to only a 34.5% change for intravenously-dosed midazolam after rifampicin treatment (Link et al. 2008). Thus, future work will include studying DDIs for drugs with both routes of administration: intravenously (i.e., liver-only system) and oral (i.e., liver and gut integrated system). Such an MPS platform would enable investigation of DDIs caused by induction at both hepatic and intestinal MPSs and evaluation of multiple PK parameters related to oral drug administration, such as clearance values in gut and liver MPS, gut MPS drug permeability and bioavailability (Tsamandouras et al. 2017). PK parameters evaluated with human MPSs may further improve the accuracy of in vitro in vivo extrapolation (IVIVE) for DDI studies by providing a human physiology-relevant experimental system. (Mehta, Karnam, and Madgula 2024)

The LTC platform was designed to address the major challenges in the MPS field for ADME applications (Fowler et al. 2020) by incorporating features such as medium recirculation, thermoplastic tissue chip material for low non-specific binding, sufficient volume for media-based sampling, and low evaporation. The bioengineered millifluidic tissue chip enabled the long-term PK assessment of both low and high clearance compounds, which is critical for assessing DDIs *in vitro*. These results demonstrated a robust and reproducible platform supporting long-term liver-mediated DDI studies. Further, this platform serves as a foundation for future development of multi-organ interconnected systems that could better predict human responses (Edington et al. 2018; Wang et al. 2019) and provide physiologically relevant data before first-in-human studies (Rajan et al. 2023).

This work was supported by the National Institute of Health National Center for Advancing Translational Sciences [NIH 1R44TR004524]; and National Institute of Health National Institute of General Medical Sciences [NIH 1R43GM143980].

## ACKNOWLEDGEMENTS

The authors would like to thank Donald Tweedie for his valuable guidance and discussions during the manuscript review process. We are also grateful to Jason Sherfey for his insightful discussions on data analysis.

## AUTHORSHIP CONTRIBUTIONS

**Participated in research design:** Ohri, Rajan, Cirit

**Conducted experiments:** Ohri, Nichols

**Performed data analysis:** Ohri, Parekh

**Wrote or contributed to the writing of the manuscript:** Ohri, Rajan, Cirit

