## Supplementary Material for "Utilization of a Human Liver Tissue Chip for Drug-Metabolizing Enzyme Induction Studies of Perpetrator and Victim Drugs"

**Affiliation:**
SO, PP, LN, SR, MC - Javelin Biotech, Inc., 299 Washington Street, Woburn, Massachusetts 01801, USA

**Bioanalysis:** HPLC grade water and HPLC grade acetonitrile (ACN) were obtained from Fisher Scientific. Formic Acid (98.0-100%) and Ammonium Formate (97%) were obtained from Sigma Aldrich, Inc. The following reference materials: 5-hydroxy omeprazole, 4-hydroxy diclofenac, hydroxy tacrine, 1-hydroxy midazolam, dextrorphan, tolbutamide, 4-hydroxy diclofenac-13C6, dextrorphan-d3, alprazolam, alprazolam-d8, rifampicin, rifampicin-d3, midazolam-d4, and midazolam and were purchased from Sigma Aldrich, Inc., Missouri, United States or Caymen Chemical, Michigan, United States.

Stock solutions of each compound were first prepared in DMSO at 1 mg/ml. Standard calibration working solutions were prepared by serial dilution in ACN over the range of 1.25 to 10,000 ng/ml. Then, 12.0 µL of the working solutions were added into 48.0 µL of blank matrix for a calibration curve range of 0.25 to 2000 ng/ml. The standards and study samples were then vortexed at 1000 rpm for 3 min. For matrix blank and control zero samples, 12.0 µL of acetonitrile was added to 48.0 µL of blank matrix. For calibration standard and QC samples, 240 µL of acetonitrile containing internal standards at 50 ng/ml each was added to all wells. Refer to Table S2 – S5 for internal standard compounds. For matrix blanks, 240 µL of pure acetonitrile was added. For study samples, 200 µL of acetonitrile containing internal standards at 50 ng/ml each was added to 50 µL of sample. The plates were sealed and vortexed at 1700 rpm for 3 min. The plates were then centrifuged at 4000 rpm for 10 min at room temperature. 100 µL of the supernatant was then transferred to a clean Axygen 96-well collection plate containing 100 µL ultrapure water. The plates were sealed and vortexed at 1700 rpm for approximately 3 min, then injected onto the LC-MS/MS system. The chromatographic system was allowed to equilibrate with each initial mobile phase condition prior to analysis.

The chromatographic system for study A was an Agilent 1290 Infinity II High-Throughput System consisting of a G7116B multicolumn thermostat, a G7167B multisampler, and a G7120A high speed pump. The chromatographic system for studies B, C, and D included a Shimadzu Nexera X2 CBM-20A communication bus module, Shimadzu Nexera X2 LC-30AD pumps, Shimadzu Nexera X2 SIL-30ACMP autosampler, and Shimadzu DGU-20A5 degassers. For studies A, B and C Mobile phase A consisted of water containing 0.1% (v:v) formic acid and mobile phase B consisted of acetonitrile containing 0.1% (v:v) formic acid. For study D, Mobile Phase A consisted of 5mM Ammonium Formate in H2O and Mobile Phase B was acetonitrile. For study A the flow rate and column temperature was 0.8 ml/min and 55°C, respectively. For studies B and C the flow rate and column temperatures were 0.8 ml/min and 50°C, respectively. For study D the flow rate and column temperatures were 1.0 ml/min and 50°C, respectively. The chromatographic separations were achieved for studies A and D using an Agilent Zorbax SB-C18, 3.5 µM, 4.6 x 50mm column, P/N 835975-902. For studies B and C an Agilent Zorbax SB-C18, 5 µM, 2.1 x 50mm column, P/N 860975-902. The gradient programs are tabulated in Table S1.

The mass spectrometric detector utilized to conduct sample analysis for study A was an Agilent 6495D Triple Quadrupole equipped with an Agilent Jet Stream™ ESI source. The source conditions are as follows: Gas Temp 250 °C, Capillary Voltage 3000 V, and Nozzle Voltage 1500 V.

The mass spectrometric detector utilized to conduct sample analysis for studies B, C, and D was a Sciex Triple Quad™ 5500+ equipped with a IonDrive™ Turbo V source. The source conditions were as follows: positive ion electrospray, CUR 35, CAD 10, IS 5500, TEM 500, GS1 50, GS2 60, EP 10, and CXP 10.

The data acquisition was performed using multiple reaction monitoring (MRM). MRM transitions can be found in Tables S2 to S5. Data acquisition and chromatographic review were performed using Agilent MassHunter, version 10.1 for study A and Applied Biosystems/MDS Sciex Analyst, version 1.7.2 for studies B, C, and D. The calibration curves utilized $1/x^2$ weighting

**Table S1 Chromatographic Gradients**

**Study A**

| Time (min) | Module | Event | Parameter |
|---|---|---|---|
| 0.2 | Pumps | Pump B Conc. | 15 |
| 1.5 | Pumps | Pump B Conc. | 95 |
| 2.25 | Pumps | Pump B Conc. | 95 |
| 2.35 | Pumps | Pump B Conc. | 15 |
| 3.00 | Controller | Stop | |

**Study B and C**

| Time (min) | Module | Event | Parameter |
|---|---|---|---|
| 0.01 | Pumps | Pump B Conc. | 20 |
| 2.00 | Pumps | Pump B Conc. | 95 |
| 2.50 | Pumps | Pump B Conc. | 95 |
| 2.60 | Pumps | Pump B Conc. | 20 |
| 3.00 | Controller | Stop | |

**Study D**

| Time (min) | Module | Event | Parameter |
|---|---|---|---|
| 0.1 | Pumps | Pump B Conc. | 30 |
| 2.00 | Pumps | Pump B Conc. | 95 |
| 2.50 | Pumps | Pump B Conc. | 95 |
| 2.60 | Pumps | Pump B Conc. | 30 |
| 3.00 | Controller | Stop | |

**Table S2. Study A MRM Parameters**

| MRM Parameters | | | | | |
|---|---|---|---|---|---|
| **Q1** | **Q3** | **Dwell time** | **Compound** | **DP** | **CE** |
| **Analytes** | | | | | |
| 362.1 | 214 | 200 | 5-hydroxy omeprazole[1] | N/A | 8 |
| 342.1 | 323.9 | 200 | 1-hydroxy midazolam[1] | N/A | 26 |
| 215.1 | 197 | 200 | hydroxy tacrine[1] | N/A | 37 |
| 312 | 230 | 200 | 4-hydroxy diclofenac[2] | N/A | 37 |
| 258.2 | 157.2 | 200 | dextrorphan[3] | N/A | 75 |
| **Internal Standards** | | | | | |
| 271.0 | 91.1 | 200 | Tolbutamide | N/A | 33 |
| 318.1 | 236.9 | 200 | 4-hydroxy diclofenac-13C6 | N/A | 27 |
| 261.2 | 133.2 | 200 | dextrorphan-d3 | N/A | 55 |

[1]Internal standard used was Tolbutamide.

[2]Internal standard used was 4-hydroxy diclofenac-13C6.

[3]Internal standard used was dextrorphan-d3.

**Table S3. Study B MRM Parameters**

| MRM Parameters | | | | | |
|---|---|---|---|---|---|
| **Q1** | **Q3** | **Dwell time** | **Compound** | **DP** | **CE** |
| **Analyte** | | | | | |
| 309 | 281.3 | 150 | alprazolam | 132 | 38 |
| **Internal Standard** | | | | | |
| 317 | 289.3 | 50 | alprazolam-d8 | 132 | 60 |

**Table S4. Studies C MRM Parameters**

| MRM Parameters | | | | | |
|---|---|---|---|---|---|
| **Q1** | **Q3** | **Dwell time** | **Compound** | **DP** | **CE** |
| **Analyte** | | | | | |
| 823.4 | 791.6 | 150 | rifampin | 143 | 27 |
| **Internal Standard** | | | | | |
| 826.4 | 794.6 | 50 | rifampin-d3 | 143 | 44 |

**Table S5. Study D MRM Parameters**

| MRM Parameters | | | | | |
|---|---|---|---|---|---|
| **Q1** | **Q3** | **Dwell time** | **Compound** | **DP** | **CE** |
| **Analyte** | | | | | |
| 342.1 | 323.9 | 150 | midazolam | 90 | 52 |
| **Internal Standard** | | | | | |
| 346.2 | 328.3 | 50 | midazolam-d4 | 90 | 40 |

**Table S6**

Donor demographics of all the cryopreserved primary human hepatocytes (PHH) used for the culture of liver MPS on the LTC.

| Vendor | Identifier | Sex | Age | Ethnicity | Cause of death |
|---|---|---|---|---|---|
| LifeNet Health | 2214423 | Female | 28 | Caucasian | Anoxia |
| AnaBios Corporation | 1045 | Female | 2 | Caucasian | Anoxia/Drowning |
| ThermoFisher Scientific | Hu8339 | Female | 31 | African American | Asphyxiation |

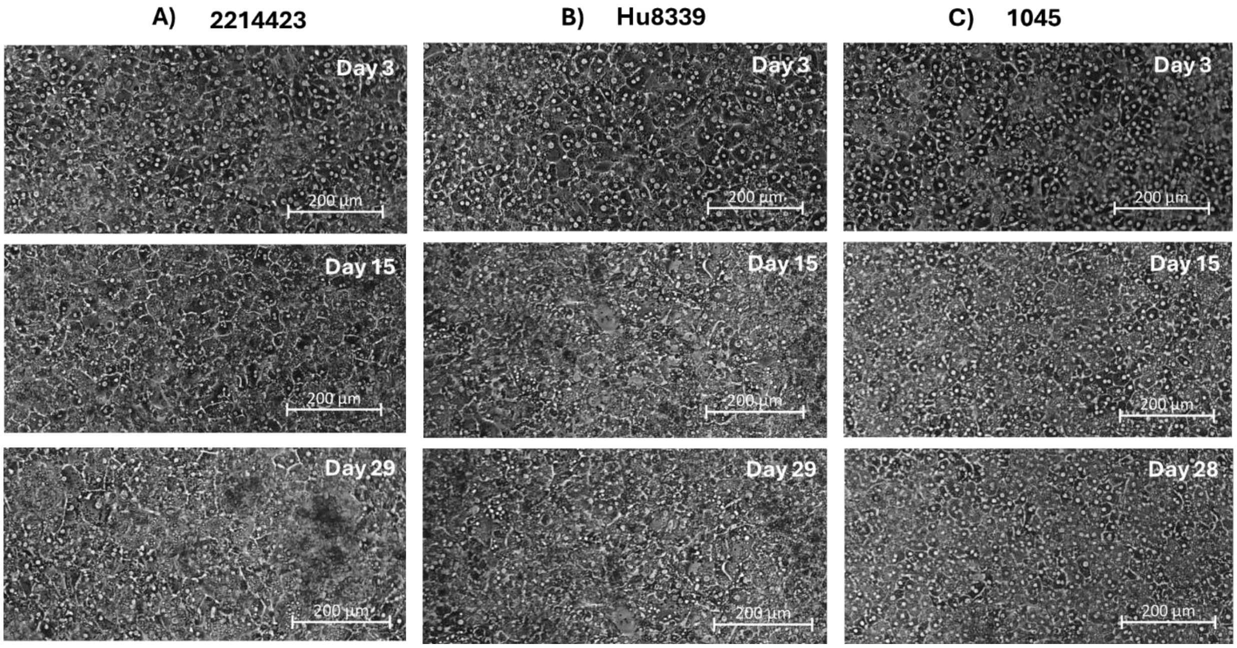

**Figure S1:** Long-term morphological characterization of liver MPS from three different donors. Brightfield images of PHH from (A) 2214423 (B) Hu8339 (C) 1045 showing characteristic multi-nucleated and cobblestone tissue morphology maintained for over four-weeks.

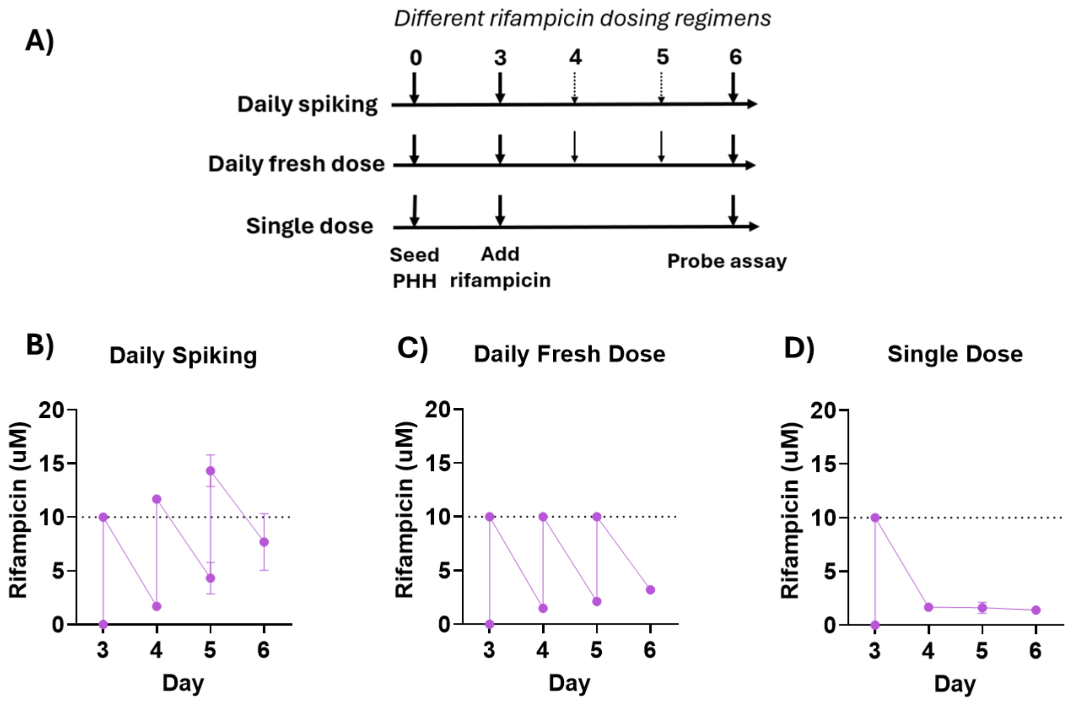

**Figure S2:** The Liver MPS (PHH Donor: 1045) was dosed with rifampicin for 72 hours to investigate the effects of three rifampicin dosing regimens - i) daily spiking, ii) daily fresh dose, and iii) single dose. (A) On day 3, using the complete medium change protocol, 1.8mL of fresh medium containing 10 µM rifampicin was added for all three-dosing regimens. For the daily spiking regimen, the medium in the sampling chamber was spiked with 1.8 µL of 10 mM rifampicin stock every 24 hours after initial dose for an effective concentration of at least 10 µM. In the daily fresh dose condition, spent rifampicin-dosed medium was replaced every 24 hours with fresh medium containing 10 µM rifampicin until 72 hrs. For the single dose regimen, no drug spiking or medium replacement was conducted throughout the 72 hours of induction pre-treatment. For each dosing regimen, the rifampicin concentration in LTC was quantified daily during the induction period between days 4 and 6 and concentration-time curve was plotted for (B) Daily spiking (C) Daily fresh dose and (D) Single dose after assuming an experimental starting rifampicin concentration of 10 µM after every fresh medium replacement and instant mixing of the spiked drug stock for an effective concentration of at least 10 µM in the LTC medium for daily spiking. Rifampicin exposure was calculated by quantifying the area-under-the-curve (AUC) of concentration over 72 hours of induction pre-treatment. Plotted data represents mean + SD of 3-4 biological replicates.

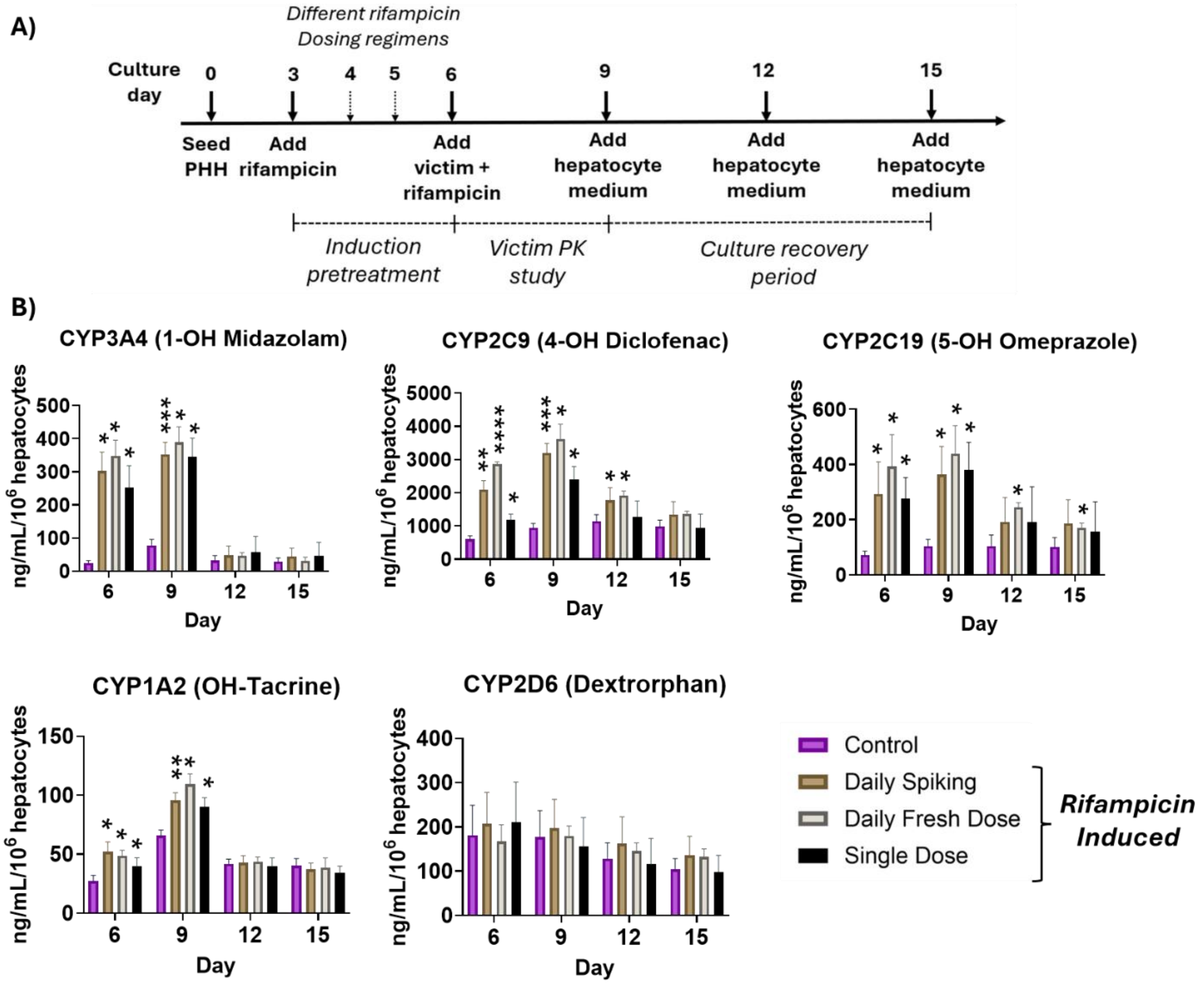

**Figure S3:** (A) The liver MPS (PHH Donor: 1045) was induced with rifampicin starting at day 3 for 72 hours with different rifampicin dosing regimens and at day 6, a single dose of 10 µM Rifampicin was added for another 72 hours (victim PK study period). On day 9, rifampicin-free hepatocyte maintenance medium was added for an additional 6 days to observe the baseline recovery of CYP activity levels. CYP activity levels were assessed on days 6, 9, 12, and 15. (B) Comparison of time course and metabolic activity recovery levels of liver MPS across different dosing regimens revealed single dose of rifampicin during induction pre-treatment resulting in similar de-induction of CYP3A4 and CYP2C19 activity as was seen with daily spiking of rifampicin (return to baseline activity levels within 3 days), however, CYP2C9 activity returned to baseline levels within 3 days as opposed to within 6 days after daily spiking of rifampicin. Daily fresh dose of rifampicin also resulted in similar CYP3A4 activity recovery compared to rifampicin daily spiking (return to baseline levels within 3 days), however, CYP2C9 and CYP2C19 activity did not completely return to the baseline level after induction pre-treatment. Statistical significance is displayed relative to control determined using un-equal variance t-test (* $.001 < p < 0.05$; ** $.0001 < p < .001$; *** $.00001 < p < .0001$; **** $p < .00001$; no significance (ns) $p > 0.05$, not shown on graph). Plotted data represents mean + SD of 3-4 biological replicates.

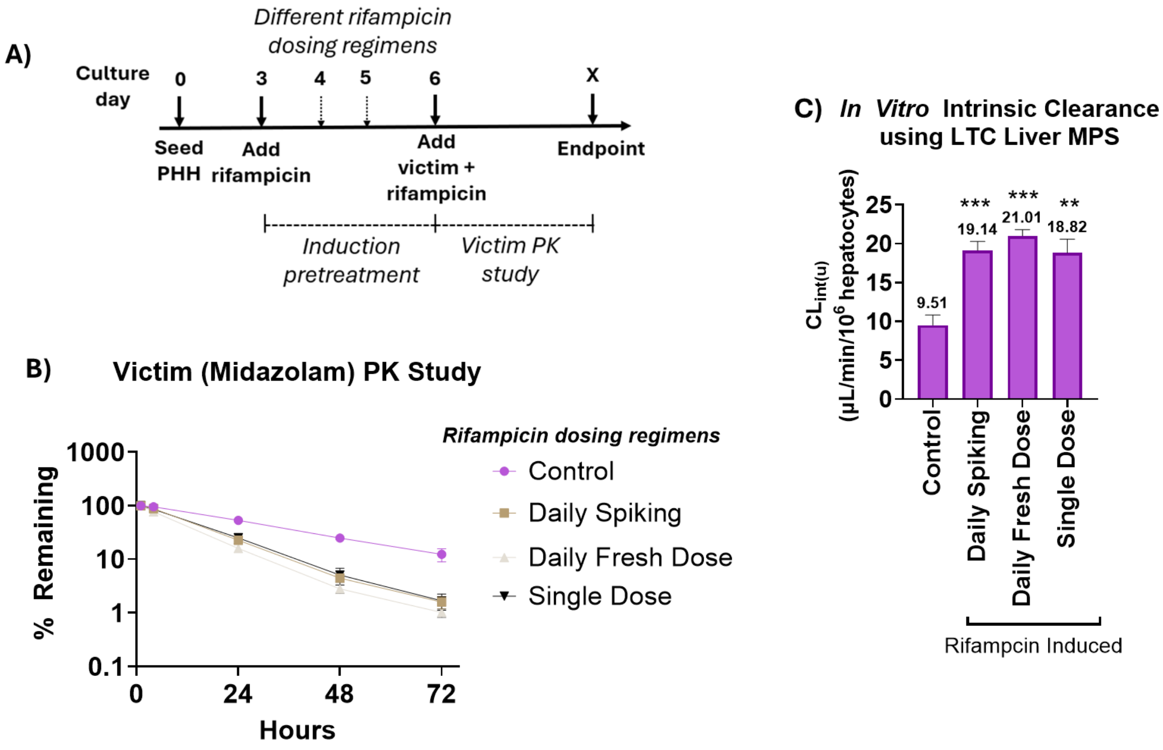

**Figure S4:** (A) The liver MPS (PHH Donor: 1045) was induced with rifampicin starting at day 3 for 72 hours using different rifampicin dosing regimens and then, at day 9, 1 µM victim drug, midazolam, was co-dosed with 10 µM rifampicin, the perpetrator drug, for the victim PK study. (B) Comparison of midazolam depletion profile after rifampicin-mediated induction with different dose regimens revealed 43-, 54-, and 42% reduction in $AUC_{0\text{-}72\ hr}$ for daily spiking, daily fresh dose, and single dose respectively, when compared to the vehicle control. (C) Unbound in vitro intrinsic clearance values revealed a 2-, 2.2-, and 1.98-fold increase in midazolam clearance after liver MPS induction with daily spiking, daily fresh dose, and single dose of rifampicin, respectively compared to the control. Statistical significance is displayed relative to control determined using un-equal variance t-test (* .001 < p < 0.05; ** .0001 < p < .001; *** .00001 < p < .0001; **** p < .00001; no significance (ns) p > 0.05). Plotted data represents mean + SD of 3-4 biological replicates.

**Table S7**

Predicted Human Hepatic Clearance ($CL_h$) Values Using Well Stirred (WS) and Parallel Tube (PT) Model

Unbound *in vitro* intrinsic clearances ($CL_{int(u)}$) of the victim drug (midazolam) was calculated for vehicle control and rifampicin induced liver MPS with different dosing (daily spiking, daily fresh dose, single dose) using the LTC substrate depletion data which was scaled to predicted human hepatic clearance values using human hepatocellularity and WS, PT models.

| Victim Drug | Condition | $CL_{int(u)}$ [µL/min/$10^6$ hepatocytes] | $CL_h$ (WS) [mL/min/kg] | $CL_h$ (PT) [mL/min/kg] |
|---|---|---|---|---|
| Midazolam | Control | 9.51 | 2.34 | 2.47 |
| | Daily Spiking | 19.14 | 4.23 | 4.67 |
| | Daily Fresh Dose | 21.01 | 4.55 | 5.07 |
| | Single Dose | 18.82 | 4.17 | 4.60 |
